# Effectors with different gears: divergence of *Ustilago maydis* effector genes is associated with their temporal expression pattern during plant infection

**DOI:** 10.1101/2020.12.10.419192

**Authors:** Jasper R.L. Depotter, Weiliang Zuo, Maike Hansen, Boqi Zhang, Mingliang Xu, Gunther Doehlemann

## Abstract

Plant pathogens secrete a variety of effector proteins that enable host colonization but are also typical pathogen detection targets for the host immune system. Consequently, effector genes encounter high selection pressures, which typically makes them fast evolving. The corn smut pathogen *Ustilago maydis* has an effector gene repertoire with a dynamic expression across the different disease stages. We determined the amino acid divergence of *U. maydis* effector candidates with *Sporisorium reilianum* orthologs, a close relative of *U. maydis*. Intriguingly, there are two distinct groups of effector candidates, ones with a respective conserved and diverged protein evolution. Conservatively evolving effector genes especially have their peak expression during the (pre-)penetration stages of the disease cycle. In contrast, expression of divergently evolving effector genes generally peaks during fungal proliferation within the host. To test if this interspecific effector diversity corresponds to intraspecific diversity, we sampled and sequenced a diverse collection of *U. maydis* strains from the most important maize breeding and production regions in China. Effector candidates with a diverged interspecific evolution had more intraspecific amino acid variation than candidates with a conserved evolution. In conclusion, we highlight diversity in evolution within the *U. maydis* effector repertoire with dynamically and conservatively evolving members.

## Introduction

Plant pathogens engage in intimate interactions with their hosts. They colonize host tissue and use host metabolites for their own benefit, which negatively impacts the host fitness. Consequently, hosts and pathogens evolve in an evolutionary arms race in which both aim to outmarch each other [1]. Plants therefore evolved an immune system to detect pathogen evasion, which then triggers the production of metabolites that prevent pathogen ingress [2]. Pathogens, on the other side, produce a range of secreted proteins that aim to circumvent the host’s immune system and also alters host metabolic processes to their own benefit [3]. These proteins are called effector proteins and are essential for host colonization, but conversely are also typical targets that are recognized by the host immune system leading to a halt of pathogen ingress. Hence, effectors can also be referred to as avirulence (Avr) effectors [4]. This selection pressure makes effectors a dynamic group of proteins that evolve fast to overcome immunity and to guarantee continued host symbiosis [5].

Although effector proteins are often collectively considered as rapid evolving, there are differences in evolutionary speed between individuals [6]. Certain effectors are indispensable, as their absence would make host colonization impossible. For instance, Pep1 is an effector of the maize smut pathogen *Ustilago maydis* that is indispensable for penetration of the host epidermis [7]. In contrast, there are effectors with accessory functions that optimize host colonization but are not necessary for pathogens to complete their life cycle. *U. maydis* contains such accessory gene clusters [8], of which the biggest one, so-called 19A cluster, contains 24 genes encoding secreted effectors [9]. Although deletion of the 19A cluster severely reduces virulence, *U. maydis* is still able to complete its life cycle. Effectors could also have overlapping functions with other effectors, which makes them redundant. To prove the redundancy of effectors is difficult as one cannot exclude that an effector contributes to colonization in particular circumstances or on particular host genotypes that have not been tested [6]. In the rice blast fungus *Magnaporthe oryzae*, one out of 78 single effector gene mutants displayed a reduction in virulence, indicating that functional redundancy occurs [10]. Furthermore, effectors can contribute to host colonization in lineage specific manner. For instance, the *U. maydis* effector Apathogenic in B73 (ApB73) is necessary for the infection of maize line B73, whereas the maize line Early Golden Bantam could be infected, although with lower virulence, when *apB73* was deleted [11]. Similarly, *U. maydis* effector UMAG_02297 contributes to full virulence in maize line CML322 but not in Early Golden Bantam [12]. Conceivably, differences in nature and function of effectors in the pathogen life cycle must impact their mutability. Following this train of thought, indispensable effectors are more likely to be conserved within populations, whereas effectors with more accessory functions can display more variations [6].

Smuts are a group of basidiomycete plant pathogens that mainly infect Poaceae [13]. In their life cycle, smuts encounter a haploid stage where they live outside the plant as a yeast form. Alternatively, smuts can live as a diploid filament inside the plant after mating of two yeasts with compatible mating tape. *U. maydis* and *Sporisorium reilianum* are two closely related smuts that both infect maize. Most smuts, including *S. reilianum*, systemically colonize their host to then produce their spores in the host influorescence [13,14]. In contrast, maize plants infected by *U. maydis* develop tumours, where teliospores are eventually formed [15]. Tumours with mature spores can already be observed in less than two weeks after infection [16]. In this time, *U. maydis* undergoes different stages during its biotrophic life cycle, in which it encounters various transcriptional changes [16]. The secretome of *U. maydis* is estimated to comprise of 467-729 genes depending on the possibility of nonconventional secretion is included and the stringency in cut-offs used for an extracellular prediction [17,18]. Secreted proteins alter their expression in a wave like fashion during the course of infection [16]. Thus, many secreted effectors must perform specific roles during infection that are especially required in a particular stage of host colonization. For instance, the *U. maydis tin2* encodes a virulence factor that stabilizes the cytoplasmic maize protein kinase ZmTTK1 leading to anthocyanin production [19]. The expression of *tin2* shows a wave-like fashion which gradually increases during infection and peaks at 6 days post infection (dpi) to then drop again at 8 dpi.

Although *U. maydis* and *S. reilianum* infect the same host, symptom development and how they colonize their host are distinct. We hypothesized that effectors which are active at different points of the infection cycle may evolve distinctly from each other and thus we investigated interspecific and intraspecific evolution of the *U. maydis* effector repertoire. Since pathogen populations encounter a variety of cultivars in cropping systems with location specific characteristics, we sampled *U. maydis* isolates from maize fields in throughout China. We sequenced representative strains of this population to search for differences in effector variation that reflect local adaptations.

## Materials and Methods

### 2.1. Effector candidate determination

The reference genome annotation of *U. maydis* was used [8]. To determine secreted proteins, the presence of signal peptides was predicted for the *U. maydis* proteins with SignalP (v5.0) [20]. Additionally, proteins with localization prediction to the plant apoplast was predicted with ApoplastP (v1.0) [21]. Transmembrane helices in proteins were predicted with TMHMM2.0c [22]. For proteins with a signal peptide, the signal peptide was excluded for transmembrane helix prediction. Proteins were also annotated for a CAZyme function with the dbCAN2 web server [23]. dbCAN2 uses three tools using HMMER, DIAMOND and a Hotpep search. If any of these three tools predicted a CAZyme function, the protein was assigned as a CAZyme. Previously published *U. maydis* gene expression data was analysed with samples from *U. maydis* grown in axenic culture and in different plant associated life cycle stages at 0.5, 1, 2, 4, 6, 8 and 12 dpi [16]. To this end, reads were filtered using the Trinity software (v2.9.1) option trimmomatic under the standard settings [24]. The reads were then mapped to the reference genome using Bowtie 2 (v2.3.5.1) with the first 15 nucleotides on the 5’-end of the reads being trimmed [25]. The reference genome was the genome assembly of *U. maydis* [8] combined with the assembly of *Z. mays* B73 version 3 [26]. Reads were counted to the *U. maydis* coding regions of genes using the R package Rsubread (v1.34.7) [27]. Significant differential expression of a locus was determined using the R package edgeR (v3.26.8) [28]. Significance of differential expression was calculated using t-tests relative to a threshold of log2 fold change of 1 with Benjamini-Hochberg correction. The cut-off of 0.05 was used.

### 2.2. *Effector divergence between* U. maydis *and* S. reilianum

One-to-one orthologs between *U. maydis* and *S. reilianum* were determined based on phylogeny using the OrthoFinder (v2.4.0) software [29,30]. To this end, protein annotations of *U. maydis* [8], *S. reilianum* [31], *Sporisorium scitamineum* [18], *Ustilago bromivora* [32], *Ustilago hordei* [33] and *Anthracocystis flocculosa* [34] were used. The amino acid sequence identity was calculated between one-to-one *U. maydis*/*S. reilianum* orthologs with more than 40% overlap using the output of blastp (v2.2.31+) [35]. The identity distribution of these orthologous was constructed using Gaussian Kernel Density Estimation with a kernel bandwidth of 4. To determine the peak expression of the effector candidate, the highest Transcripts Per Million (TPMmax) was determined for the plant associated growth stages of samples taken at 0.5, 1, 2, 4, 6, 8 and 12 dpi [16].

### 2.3 U. maydis *sequencing and variation analysis*

Ear tumour tissues were collected from different provinces in China during the summer of 2018 summer (Figure 2A). Teliospores of the *U. maydis* were isolated from plant material and germinated as previously described [36]. Single colonies were grown on PD-agar plate or YEPS light (0.4% yeast extract, 0.4% peptone, and 2% saccharose) liquid medium with 200 rpm shaking at 28°C. For DNA extraction, cells were grown overnight in YESP light at 28°C. DNA of this fungal material was extracted [37], then purified by epicentre MasterPure DNA Purification Kit (Illumina, Madison, Wisconsin, USA) according to the manufacturer’s instructions. DNA concentration was measured by Qubit Fluorometric (Waltham, Thermo Fisher, Massachusetts, USA). DNA was then sent to for library preparation and sequencing to Novogene (Beijing, China). Sequencing library preparation was done using the NEB Next® Ultra™ RNA Library Prep Kit (NEB, Ipswich, USA). Libraries (350 bp insert size) were loaded on Illumina NovaSeq6000 System for 150bp paired-end sequencing using a S4 flowcell.

**Figure 1.**
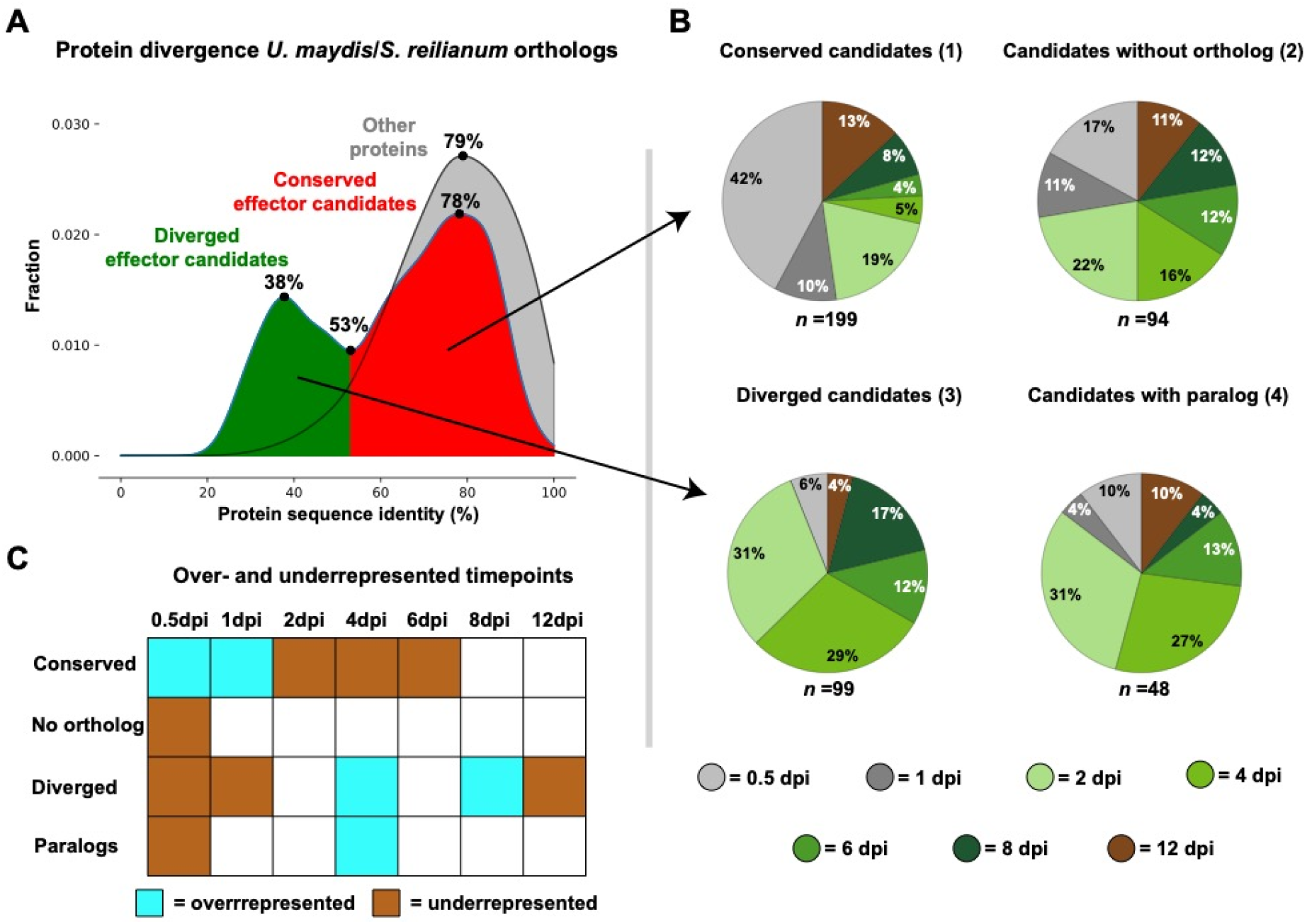
Effector candidate evolution in relation to maximum relative expression. (**A**) Protein sequence identity distribution between *U. maydis* and *S. reilianum* one-to-one orthologs was determined. The minimum and maxima in the distributions are indicated. (**B**) The distribution of the maximum relative expression point for *U. maydis* effector candidates was determined for (1) ones with a one-to-one ortholog ≥ 53% protein sequence identity, (2) ones without an ortholog, (3) ones with a one-to-one ortholog < 53% protein sequence identity and (4) ones with either *U. maydis* or *S. reilianum* containing paralogs. (**C**) Over- and underrepresentation of maximum relative expression points in the effector candidate categories were calculated with a two-sided Fisher exact test. Significance was considered for *p*-values <0.05.

**Figure 2.**
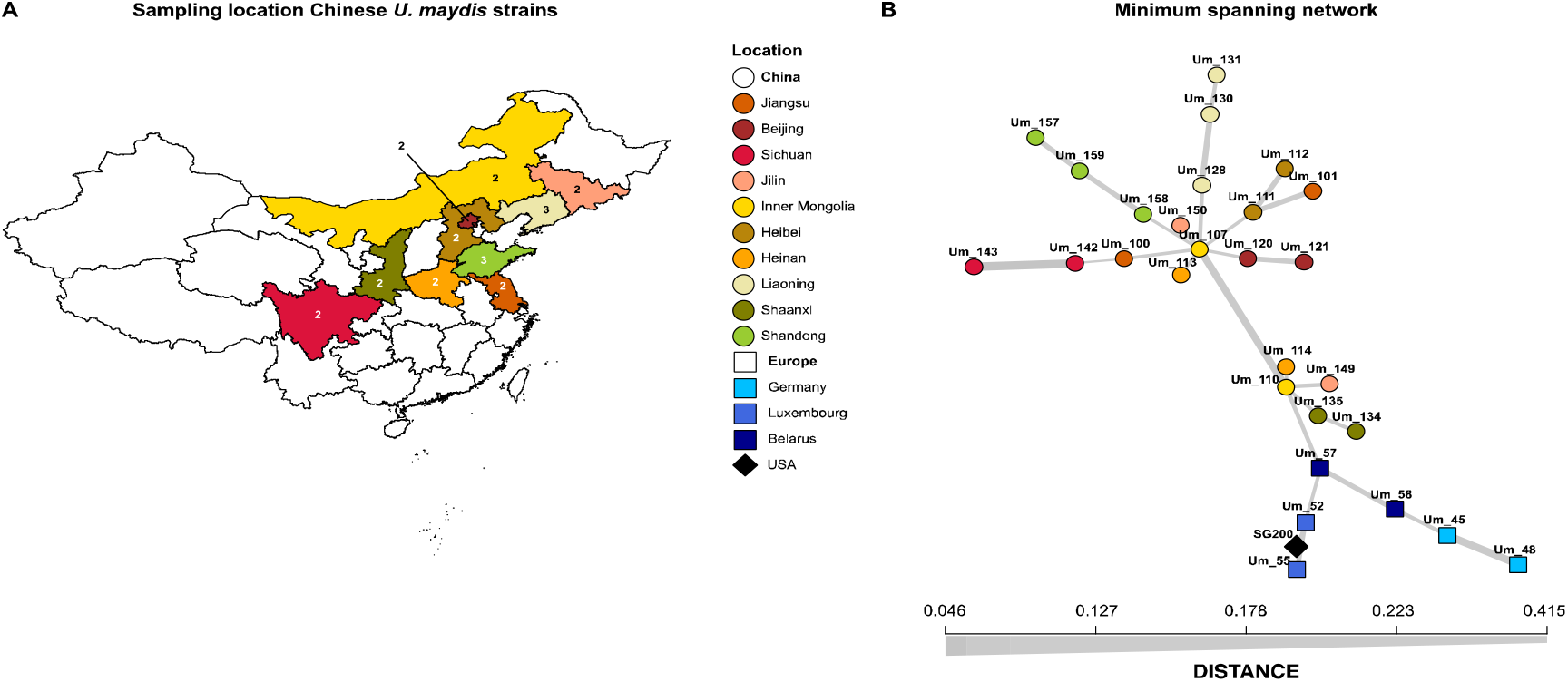
Geographic origin and genetic genealogy of *Ustilago maydis* strains. (**A**) Geographic origin of Chinese *U. maydis* isolates. The numbers associated to the provinces indicate the number of isolates that were selected for whole genome sequencing. (**B**) Minimum spanning network of the sequenced *U. maydis* population. Every isolate represents one circle/square/diamond in the figure. The isolates are connected to each other with lines, which thickness is related to the Hamming distance between the isolates.

Paired-end *U. maydis* reads were filtered using Trimmomatic v0.39 with the settings “LEADING:3 TRAILING:3 SLIDINGWINDOW:4:15 MINLEN:100”, only reads that remained paired after filtering were used for read mapping [38]. The *U. maydis* reference genome of strain 521 was used as a reference for read mapping [8]. Reads were mapped to the reference genome using BWA-MEM (v.0.7.17) [39]. Duplicated reads were tagged and then variants were called with GATK after filtering using the following requirements: SNP = “QD < 2.0 ‖ FS>60.0 ‖ MQ < 40.0” and INDEL = “QD < 2.0 ‖ FS>200.0”. Minimum spanning network of the *U. maydis* isolates was produced with Poppr (v.2.8.6) [40]. The called variants were implemented in the gene coding sequences using bcftools (v.1.9). Bcftools concat (--remove-duplicates) was used to combine the filtered SNP and INDEL Variant Call Format (VCF) files. Normalization with bcftools norm and filtering with bcftools filter (--IndelGap 5) were performed before the coding sequences were altered.

Variants were determined for the genes of the *U. maydis* strain 521 reference annotation [8]. For absence/presence polymorphisms, a homolog was considered absent in one of the *U. maydis* strains (1) if the coding sequence did not contain a start codon, (2) if less than 60% of the gene locus had reads mapped to it and (3) if a mutation resulted in a premature stop codon that reduced the protein length to at least 60% of the reference. Length polymorphisms were considered any mutation that led to a change in encoded protein length that is no smaller than 60% of the length of the reference protein. For the protein variation, pair-wise comparisons, including all the newly sequenced *U. maydis* strains and the strain 521 reference genome, were performed. To this end, all alleles in the population were aligned for every *U. maydis* genes using MAFFT (v7.464) option “--auto” [41]. Consequently, the dn was calculated for every strain pair with SNAP [42]. The dn variance between these pair-wise comparisons was then calculated. Genes that contain absence/presence polymorphisms or had a frame shift in one of the strains were excluded from this analysis.

### 2.4. Data availability

The reads of the newly sequenced *U. maydis* strains are publicly available in the BioProject PRJNA674756.

## Results

### U. maydis *effector candidates diverge with two different speeds*

To study the evolution of effectors, candidate genes have to be determined. Proteins were defined as effector candidates if a signal peptide or an apoplast localization was predicted. Furthermore, effector candidates did not contain predicted transmembrane helices or a carbohydrate active enzyme (CAZyme) function. Finally, *U. maydis* effector candidate genes were upregulated in at least one of the plant associated growth stages of samples taken at 0.5, 1, 2, 4, 6, 8 and 12 dpi in comparison to growth in axenic culture [16]. Using this methodology, 440 effector candidates were predicted. One-to-one orthologs of these candidates with *S. reilianum* were determined based on phylogeny. In total, 298 one-to-one orthologs were determined, whereas 94 effector candidates did not have an ortholog (Table S1). The remaining 48 effector candidates did have a *S. reilianum* ortholog but were excluded because they had paralogs *in U. maydis* and/or *S. reilianum*. To study effector evolution in the *U. maydis*/*S. reilianum* divergence, the protein sequence identity of the one-to-one orthologs was determined (Table S1). Intriguingly, the protein identity had a bimodal distribution with a minimum at 53% and maxima at 38 and 78% (Figure 1A). In total, 199 effector candidates had a one-to-one ortholog with at least 53% sequence identity (conserved effector candidates hereafter), whereas 97 had a sequence identity lower than 53% (diverged effector candidates hereafter) (Table S1). In contrast, the protein sequence identity of other *U. maydis* genes that are not effector candidates had a unimodal distribution with their one-to-one *S. reilianum* orthologs with a maximum at 79% (Figure 1A). Thus, in the *U. maydis*/*S. reilianum* divergence, effector protein sequences evolved at two different speeds. The conserved group evolved protein sequence alterations at a similar pace as non-effector candidates, whereas the effectors of the diverged group encountered more changes.

### Effector gene divergence is associated with their temporal expression pattern

To see if sequence divergence is linked to gene expression, the plant associated time point with the highest Transcripts Per Million (TPMmax) was determined. Data for seven time points were available across the *U. maydis* disease cycle [16]. Samples at 0.5 and 1 dpi represent the (pre-)penetration stages of the disease cycle. At 2, 4 and 6 dpi *U. maydis* colonizes and proliferates within the host. At 8 dpi, fungal hyphae cluster in tumour tissue, whereas at 12 dpi mature spores are formed. The TPMmax was the most frequently present at 0.5 dpi (25%) followed by 2 dpi (24%), 4 dpi (15%), 8 dpi (10%), 12 dpi (10%), 6 dpi (8%) and 1 dpi (7%) (Table S1). Effectors with a TPMmax at 0.5 and 1 dpi were overrepresented in the group of conserved effectors, whereas ones with a TPMmax in 2, 4, and 6 dpi were underrepresented (Figure 1B-C, Fisher exact test, *p*-value < 0.05). Conversely, diverged effecters are underrepresented in effectors with an early TPMmax (0.5 and 1 dpi), whereas ones with TPMmax at 4 and 8 dpi are overrepresented (Figure 1B-C). Also, effectors with a TPMmax at 12 dpi, when mature spores are formed, were underrepresented in the diverged effectors. Furthermore, effectors with paralogs are enriched for members with a 4 dpi TPMmax and underrepresented in the 0.5 dpi TPMmax groups (Figure 1B-C). *U. maydis* effector genes without an ortholog in *S. reilianum* have the most even distribution of TPMmaxes, as only timepoint 0.5 dpi is significantly underrepresented (Figure 1B-C). In conclusion, genes encoding effector candidates with a conserved evolution have an early peak expression, and therefore are probably especially important in the onset of the disease cycle, i.e. during filament formation and host penetration. In contrast, genes coding for effector candidates with a more dynamic evolution through sequence alteration and duplication play a prominent role during fungal proliferation within the host. Remarkably, effectors with a previously tested organ-specific function (7 out of 7) or cell-type specific expression (12 out of 16) group into the dynamic evolving effectors (Table 1) [15,43].

**Table 1.** Features of *Ustilago maydis* effectors. TPMmax is the plant associated time point with the highest Transcripts Per Million.

| Effector | Gene number | TPMmax | Effector category* | Reference |
| --- | --- | --- | --- | --- |
| **Functionally characterized genes** |  |  |  |  |
| Rsp3 | UMAG_03274 | 2 dpi | no ortholog | [44] |
| Pit2 | UMAG_01375 | 4 dpi | diverged | [45] |
| Pep1 | UMAG_01987 | 2 dpi | conserved | [7] |
| Tin2 | UMAG_05302 | 2 dpi | diverged | [19] |
| Cmu1 | UMAG_05731 | 4 dpi | diverged | [46] |
| See1 | UMAG_02239 | 2 dpi | diverged | [47] |
| **Organ-specific effectors** |  |  |  | [43] |
| / | UMAG_05318 | 4 dpi | paralog | / |
| See1 | UMAG_02239 | 2 dpi | diverged | [47] |
| / | UMAG_11060 | 2 dpi | diverged | / |
| / | UMAG_01690 | 2 dpi | diverged | / |
| / | UMAG_05306 | 4 dpi | diverged | / |
| / | UMAG_05311 | 4 dpi | no ortholog | / |
| / | UMAG_03650 | 6 dpi | diverged | / |
| **Effector candidates with cell-type specific expression** |  |  |  | [15] |
| / | UMAG_05312 | 8 dpi | diverged | / |
| / | UMAG_00628 | 2 dpi | diverged | / |
| / | UMAG_00904 | 2 dpi | conserved | / |
| / | UMAG_02381 | 12 dpi | conserved | / |
| / | UMAG_05222 | 0.5 dpi | conserved | / |
| / | UMAG_00793 | 2 dpi | paralog | / |
| / | UMAG_02473 | 2 dpi | diverged | / |
| / | UMAG_05318 | 4 dpi | paralog | / |
| / | UMAG_03024 | 0.5 dpi | conserved | / |
| / | UMAG_02540 | 2 dpi | diverged | / |
| / | UMAG_03232 | 8 dpi | paralog | / |
| / | UMAG_03746 | 8 dpi | diverged | / |
| / | UMAG_05319 | 4 dpi | diverged | / |
| / | UMAG_05926 | 6 dpi | diverged | / |
| / | UMAG_02294 | 8 dpi | diverged | / |
| / | UMAG_03650 | 6 dpi | diverged | / |

\*Four effector categories are described:
- (1) “conserved” effector candidates have a one-to-one *Sporisorium reilianum* ortholog with a conserved protein sequence evolution - (2) “no ortholog” effector candidates do not have a *S. reilianum* ortholog - (3) “diverged” effector candidates have a one-to-one *S. reilianum* ortholog with a diverged protein sequence evolution - (4) “paralog” effector candidates have *U. maydis* and/or *S. reilianum* paralogs

### High interspecific divergence corresponds to higher intraspecific diversity

To see if differences in effector candidate evolution are also associated with intraspecific variation, we collected common smut tumour tissues from China and isolated *U. maydis* strains. Galls were collected from maize growing field under natural disease condition from 10 different provinces, which represented the main maize breeding and production regions in China (Figure 2A). In total, 28 *U. maydis* strains were sequenced, including 22 strains from the 10 maize growth regions in China, as well as 6 strains from 3 different European countries (Figure 2, Table S2). On average, 1.31 GB of 150bp paired-end reads were per strain obtained through the Illumina NovaSeq6000 System. The previously sequenced *U. maydis* strain SG200 was also included in the analysis, which is a solopathogenic lab strain derived from the wild type strain FB1 [8,48]. Strain FB1 is a cross between *U. maydis* strains 521 and 518, which are both originate Minnesota, USA [8,49]. Reads were mapped to the *U. maydis* strain 521 reference genome and variants were determined [8]. In total, 107,512 single-nucleotide polymorphisms (SNPs) were called and used to construct a minimum spanning network (Figure 2B). Here, the Chinese isolates clustered together, and the European isolates clustered with SG200, indicating the Chines population is distinct from the European and American ones.

Determined variants were used to analyse the intraspecific effector candidate diversity. The fraction of genes that were absent in at least one of the *U. maydis* strains was calculated. For non-effectors, 6.4% of the genes are absent in at least one of the *U. maydis* strains (Figure 3A).

**Figure 3.**
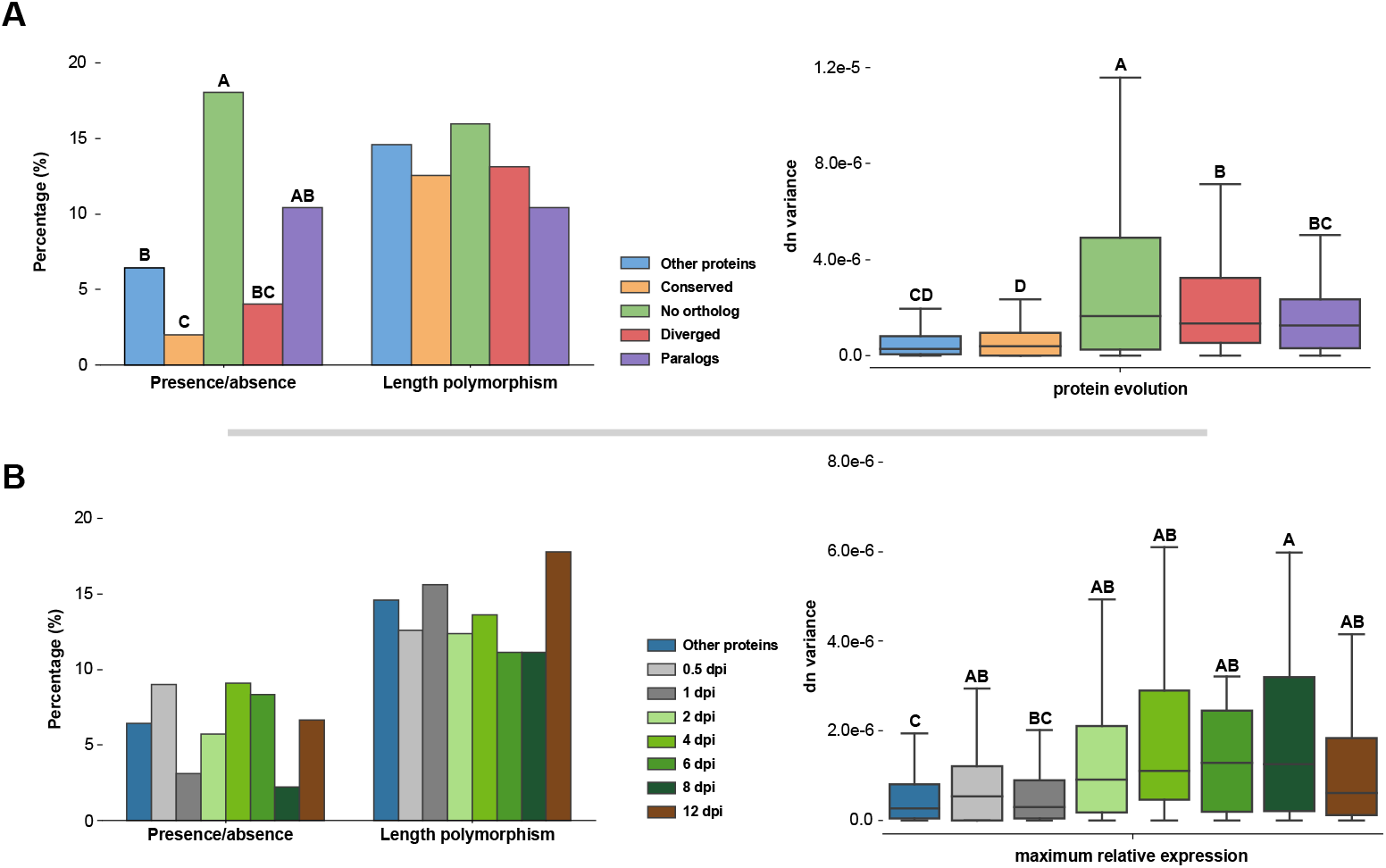
Intraspecific variation between different types of effector candidates. Differences in presence/absence polymorphisms, length polymorphisms and nonsynonymous substitutions per nonsynonymous DNA site (dn) variation between different effector candidate types were calculated. (**A**) Variation is depicted for the effector candidate categories as described in Figure 1. (**B**) Variation is depicted for effector candidates with different maximum relative expression points. Different levels of significance are indicated with A-D. In the left panels, significance was calculated with the two-sided Fisher exact test and in the right panels with the Welch’s t-test. Differences were assigned significant if *p*-value < 0.05. For plots without A-D indication, no significant differences were found.

This is a significantly larger fraction than the conserved effector candidates, which had only 2.0% of their members absent in one of the strains (Table S1, Fisher exact test, *p*-value = 0.0071). In contrast, 18.1% of the *U. maydis* effector candidates without a *S. reilianum* ortholog had absence in the population, which is a significantly higher fraction than for non-effector candidates (Table S1, Fisher exact test, *p*-value = 0.00011). Effector candidates with a diverged sequence evolution and containing paralogs had 4.0 and 10.0% of its members absent in at least one *U. maydis* strain, respectively (Figure 3A, Table S1). This was not significantly different from the non-effector candidates. In conclusion, conserved effectors candidates contain less absence/presence polymorphisms in the *U. maydis* population, whereas *U. maydis* specific effectors contain more compared to non-effector candidates. Absence of effectors in at least one of the *U. maydis* strains ranged from 2.2-9.1% for effector candidates with different TPMmax (Figure 3B, Table S1). These differences between effector TPMmax groups are not significant (Fisher exact test, *p*-value < 0.05). Furthermore, 14.6% of non-effector candidates have at least one *U. maydis* strain with a length polymorphism, which is a similar fraction to any of the effector types that ranged from 10.4 to 16.0% (Figure 3A, Table S1). Similarly, differences in length polymorphisms between effector TPMmax groups were also not significant and ranged from 11.1 to 17.8% (Figure 3B, Table S1). In conclusion, length polymorphisms occur evenly across the different effector types.

We then studied amino acid diversity of proteins within the population. To enable comparison between genes, we calculated the nonsynonymous substitutions per nonsynonymous DNA sites (dn) between every strain pair combination. For every gene, the dn variation within the population was then calculated. The median observed dn variation for non-effector candidates is 2.72e-7, which is significantly less than species specific and diverged effector candidates. (Figure 3A, Welch’s t-test, *p*-value < 0.05).

Similar to the observed effector presence/absence polymorphisms, conserved effector candidates and effector candidates without ortholog have significantly the lowest and highest dn variation with a median of 3.97e-7 and 1.64e-6, respectively (Table S1, Welch’s t-test, *p*-value < 0.05). Diverged effector candidates and effector genes with paralogues had intermediate levels of diversity with a median of 1.35e-6 and 1.27e-6, respectively (Table S1). In comparing the dn variation with effector TPMmax groups, the dn variation for non-effector candidates was not significantly different with the 1 dpi TPMmax group, whereas the other dpi groups were significantly more diverse (Figure 3B, Table S1, Welch’s t-test, *p*-value < 0.05). In conclusion, generally more intraspecific variation can be observed for effector candidates with a dynamic evolution through sequence divergence, presence/absence polymorphism or duplication, which are associated with TPMmaxes during host colonization and fungal proliferation stages of the *U. maydis* disease cycle (Figure 4). In contrast, the least intraspecific variation exists in conserved effectors that are associated with TPMmaxes in the (pre-)penetration stages.

**Figure 4.**
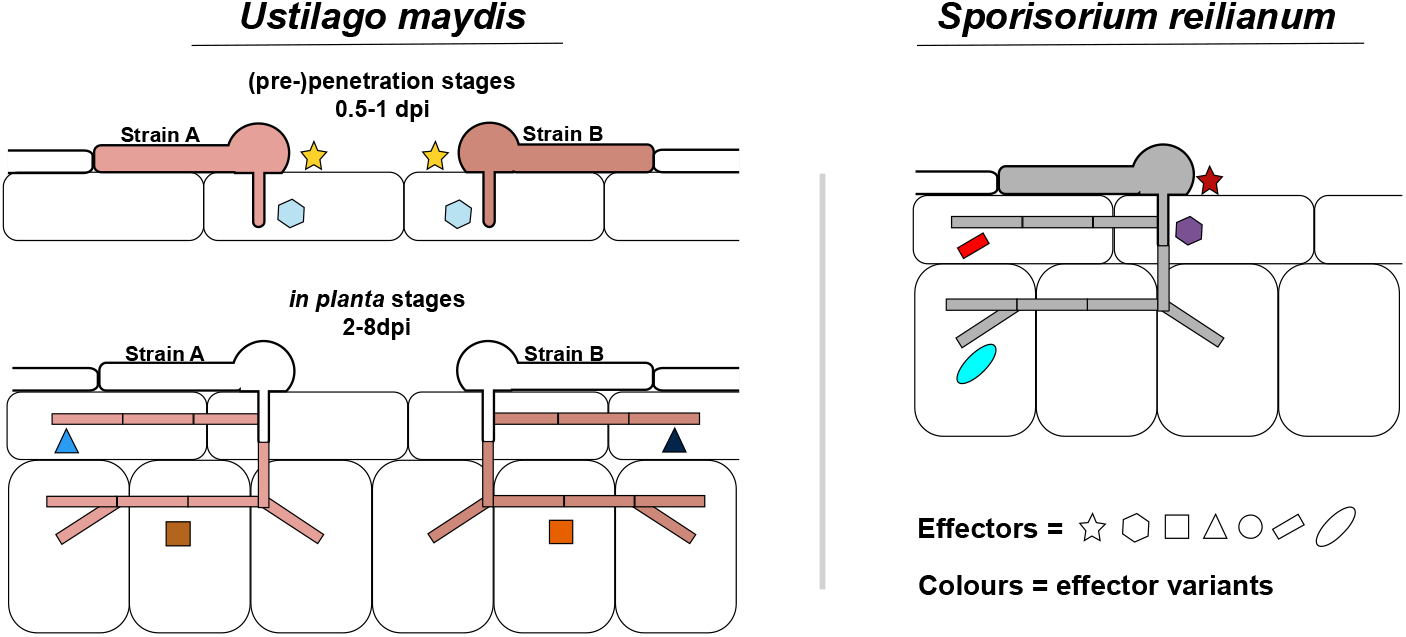
Model *Ustilago maydis* effector evolution. Effectors with a peak gene expression during the (pre-)penetration stages of the disease cycle generally encounter a more conserved evolution with less intra- and interspecific amino acid variation. In contrast, diverged effectors with higher intra- and interspecific amino acid variation generally have a peak gene expression during *in planta* colonization.

## Discussion

Effector proteins are often praised for their diversity, as the host/pathogen co-evolution fosters their innovation [50]. Indeed, many effectors lack a conserved domain and do not have homologs in other species, but conserved effectors do also exist [51–53]. The sequence identity of *U. maydis* effector candidates with one-to-one *S. reilianum* orthologs has a bimodal distribution. In total, 41% of *U. maydis* effector candidates had a one-to-one ortholog in *S. reilianum* that encountered protein divergence similar to other non-effector candidates (Figure 1A). Other effector candidates had the typical more dynamic evolution through sequence divergence, presence/absence polymorphism or duplication [18,54,55]. Intriguingly, conserved effector gene candidates are especially highly expressed during (pre-)penetration stages in contrast to candidates with a more dynamic evolution that peak at *in planta* fungal proliferation stages (Figure 1B-C). This indicates that *U. maydis* and *S. reilianum* effector repertoires mainly differentiated in effectors important for colonization inside the host, whereas effectors for the (pre-)penetration phases are more conserved. *U. maydis* effectors with a TPMmax during the (pre-)penetration phases (0.5 or 1 dpi) have hitherto not been characterized (Table 1). However, Pep1 is an effector with a conserved sequence evolution typical for the 0.5 and 1 dpi TPMmax groups but has a TPMmax at 2 dpi (Table 1). Pep1’s conserved sequence evolution corresponds to its functional conservation in various smut species [53], whereas the function of the diverged *U. maydis* effector Pit2, an apoplastic cysteine protease inhibitor, is only partially conserved in *S. reilianum* (Table 1) [45,56,57]. Although sequence conservation might go hand in hand with functional conservation, there are exceptions to this. For instance, repetitive secreted protein 3 (Rsp3) is an *U. maydis* effector protein that is required for full-virulence and anthocyanin accumulation [44]. Rsp3 has a *S. reilianum* ortholog of only 34% sequence identity but was in our analysis classified as “no ortholog”, as less than 40% of the Rsp3 sequence has coverage with its *S. reilianum* ortholog (Table 1). Despite this difference in sequence identity, the reduction in virulence in the *rsp3* mutant could be fully complemented by the *S. reilianum Rsp3* ortholog [44].

The interspecific evolutionary dynamic of *U. maydis* effectors generally corresponds to the intraspecific one as effectors with a more dynamic evolution display more absence/presence polymorphisms and protein variants than conserved effectors (Figure 3A). High intraspecific variety is especially present for *U. maydis* specific effectors. The absence of an effector in *S. reilianum* can be an indication for their accessory nature, as this close relative is able to colonize the same plant host without these effectors. Consequently, more intraspecific variation is observed as the impact of absence or amino acid alterations have a relatively lower impact of the fitness of *U. maydis* compared to effectors with indispensable functions. Alternatively, high intraspecific diversity can have a functional origin. *U. maydis* specific effectors may encounter high selection pressures that lead to a severe extent of sequence alteration making that homology to its *S. reilianum* (functional) ortholog is very low, exemplified by Rsp3 [44]. Higher variation in amino acid sequence within the *U. maydis* population shows that the outcomes of higher selection pressure can also be observed intraspecifically, as these effectors need to adapt more quickly to cultivar specific response or differences in cropping systems (Figure 3A). However, high selection pressure does not explain why *U. maydis* specific effectors encounter more absence/presence polymorphisms, indicating that a generally more accessory nature of *U. maydis* specific effectors is certainly present. To study the impact of selection pressures on the *U. maydis* population dynamics, a more extensive population sampling is needed than the one used in this study. Although there have been studies were population diversity has been assessed on a limited amount of genome loci [58], this is the first study where the intraspecific diversity of *U. maydis* has been assessed on a whole-genome scale. There is geographic segregation within the global *U. maydis* population as the Chinese strains clustered separately from the strains with a different geographic origin (Figure 2B). Interestingly, the solopathogenic strain SG200 that has an American origin clustered together with the isolates from Luxembourg, but more isolates are needed to resolve the relation between the American and European isolates. Furthermore, it would especially be interesting to perform a study on *U. maydis* population from southwest Mexico, the maize centre of origin and diversity, as the centre of diversity often correspond between pathogen and plant host [59–61].

In contrast to the *U. maydis* specific effector candidates, effector candidates with an interspecific conserved sequence evolution are hallmarked by intraspecific conservation. Theoretically, this makes conserved effector proteins ideal targets for effector recognition [6]. However, rapid recognition of conserved effectors by the host may be avoided due to a short, time point specific effector production. One of the gene expression patterns that often encode secreted proteins is the so-called red module where genes are exclusively expressed at 0.5 and 1 dpi [16]. Thus, it is plausible that a limited exposure time slows the process of host recognition down. Alternatively, as plant recognize pathogen ingress through receptors that detect patterns in the apoplast or host cytoplasm [62], it is possible that effectors secreted in the pre-penetration stage are not as well perceived by the host as ones secreted inside the host. However, during penetration, fungal plant pathogens can secrete effectors into the host from the appressorial penetration pore before host invasion [63]. Furthermore, just under half of the conserved effector candidates have a TPMmax at disease cycle stages of *in planta* fungal proliferation or sporulation (Figure 1B). Thus, as effector Pep1 exemplifies, there are effectors that target the host immune system or metabolism and evolve conservatively. As previously mentioned, these conserved effectors may be restricted in their mutability due to their indispensable function, but their recognition must then be prevented through different means [6]. For instance, recognition of conserved, indispensable effectors can be prevented through other effectors. This is illustrated by the conserved pathogen-secreted xyloglucan specific endoglucanase (PsXEG1) effector of the soybean pathogen *Phytophthora sojae* that is protected from recognition through a *PsXEG1* paralog encoding an enzymatically inactive PsXEG1 variant [64]. The enzymatically inactive variant binds more tightly than the enzymatically active variant to the soybean apoplastic glucanase inhibitor GmGIP1 that targets PsXEG1. Effectors that solely prevent recognition of other effectors evolve without other functional constraints and provide alternative targets for the host immunity to recognize instead of the (indispensable) effectors they shield.

*U. maydis* and *S. reilianum* infect the same host but diversified their colonization style in the course of evolution. Corresponding to this difference, alterations in effector repertoire were especially present in the effectors with maximum relative expression during *in planta* colonization (Figure 1B-C, 4). Alterations in effector repertoires can be episodic, which can especially be expected during drastic host changes, such as a host jump. In *U. maydis*, dynamically evolving effector candidates had higher intraspecific variation (Figure 3A, 4). This may indicate that *U. maydis* and *S. reilianum* have gradually diverged, as, through drift or selection, higher intraspecific diversity leads to the fixation of more alterations eventually leading to more diverged orthologs after speciation.

## Supplementary Materials

**Table S1**: Features of *U. maydis* effector candidates.

**Table S2:** Sequenced *Ustilago maydis* strains.

## Author Contributions

J.R.L.D., W.Z. and G.D. designed the research; J.R.L.D. performed computational data analysis, W.Z. and M.H. characterized fungal genotypes, B.Z. and M.X. performed sampling and isolation of fungal material, J.R.L.D. and W.Z. wrote the paper with input from G.D.

