## Supplementary material for "Effectors with different gears: divergence of *Ustilago maydis* effector genes is associated with their temporal expression pattern during plant infection": Table S2

**Table S2: Sequenced *Ustilago maydis* strains.**

| **Number** | **Strain name** | **Place of origin** |
| --- | --- | --- |
| 1 | Um_100 | Yangzhou, Jiangsu ,China |
| 2 | Um_101 | Yangzhou, Jiangsu ,China |
| 3 | Um_120 | Beijing, Beijing, China |
| 4 | Um_121 | Beijing, Beijing, China |
| 5 | Um_142 | Chengdu, Sichuan, China |
| 6 | Um_143 | Chengdu, Sichuan, China |
| 7 | Um_149 | Gongzhuling, Jilin, China |
| 8 | Um_150 | Gongzhuling, Jilin, China |
| 9 | Um_107 | Hohhot, Inner Mongolia, China |
| 10 | Um_110 | Hohhot, Inner Mongolia, China |
| 11 | Um_111 | Handan, Heibei, China |
| 12 | Um_112 | Handan, Heibei, China |
| 13 | Um_113 | Zhenzhou, Heinan, China |
| 14 | Um_114 | Zhenzhou, Heinan, China |
| 15 | Um_128 | Shenyang, Liaoning, China |
| 16 | Um_130 | Shenyang, Liaoning, China |
| 17 | Um_131 | Shenyang, Liaoning, China |
| 18 | Um_134 | Yangling, Shaanxi, China |
| 19 | Um_135 | Yangling, Shaanxi, China |
| 20 | Um_157 | Taian, Shandong, China |
| 21 | Um_158 | Taian, Shandong, China |
| 22 | Um_159 | Taian, Shandong, China |
| 23 | Um_45 | Germany |
| 24 | Um_48 | Germany |
| 25 | Um_52 | Luxembourg |
| 26 | Um_55 | Luxembourg |
| 27 | Um_57 | Belarus |
| 28 | Um_58 | Belarus |
